# Cross-kingdom recognition of bacterial small RNAs induces transgenerational pathogenic avoidance

**DOI:** 10.1101/697888

**Authors:** Rachel Kaletsky, Rebecca S. Moore, Lance L. Parsons, Coleen T. Murphy

## Abstract

We recently discovered that *C. elegans* can pass on a learned avoidance of pathogenic *Pseudomonas aeruginosa* (PA14) to four generations of its progeny. This transgenerational inheritance is bacterial species-specific, but how *C. elegans* recognizes and distinguishes different bacteria and transmits this information to future generations is not apparent. Here we show that small RNAs purified from pathogenic PA14 are sufficient not only to induce avoidance of pathogens in mothers, but also to confer transgenerational inheritance of this species-specific behavior for four generations, all without direct contact with pathogenic bacteria. This behavior requires the small RNA transporters SID-1 and SID-2, RNA interference pathway components, the piRNA Piwi/Argonaute pathway, a functioning germline, and TGF-β ligand *daf-7* expression in the ASI sensory neuron. Our results suggest that *C. elegans* “reads” small RNAs expressed by pathogenic bacteria, and uses this information to induce an escape behavior that lasts for four additional generations. *C. elegans* may have evolved this trans-kingdom signaling system to avoid pathogens in abundant classes of bacteria in its environment and its microbiome.

## Introduction

*C. elegans* thrive in environments replete with diverse microbial species, some of which are beneficial for the worms, while others infect and eventually kill their hosts (Dirksen et al., 2016; Samuel et al., 2016). Since *C. elegans* are bacterivores, they must distinguish beneficial and detrimental food sources to survive and reproduce. It would be logical for worms to have evolved hard-wired strategies to avoid persistent pathogens that are abundant in their environment. Alternatively, plastic learned behaviors may be necessary to appropriately respond to bacteria with dynamic levels of pathogenicity or to discriminate between pathogenic bacteria and those with closely related, non-pathogenic/nutritious species.

We recently found that a prolonged (24 h) exposure to the pathogenic bacteria *Pseudomonas aeruginosa* (PA14) induces learned avoidance not only in the mother, but also in four subsequent generations of progeny (Moore et al., 2019), before resuming its naïve level of attraction for the bacteria (Shtonda and Avery, 2006). This transient avoidance mechanism allows *C. elegans* to respond to bacteria that are abundant in the worms’ environment, but that can exist in different pathogenic states. The transgenerational inheritance of this learned avoidance behavior requires activity of the Piwi Argonaute PRG-1 and expression of the TGF-β ligand *daf-7* in the ASI neuron. Furthermore, this avoidance response is bacterial species-specific, as *C. elegans* mothers can learn to avoid the pathogen *Serratia marcesans* (Zhang et al., 2005), but do not “remember” the avoidance in subsequent generations (Moore et al., 2019). Given the specificity of this process, we wondered how *C. elegans* distinguish different bacterial pathogens, and translate this information into a long-lasting, transgenerational behavioral response.

Many animal species trigger innate and adaptive immune responses to pathogens by recognizing conserved microbial molecular motifs through toll-like receptors (TLRs)/pattern recognition receptors (PRRs) or antibodies, respectively. *C. elegans* lack an adaptive immune system, as well as many of the TLRs and PRRs present in other species (Kurz and Ewbank, 2003), and the single TLR ortholog in worms, TOL-1, does not function directly in pathogen recognition (Pujol et al., 2001). Despite the lack of evidence supporting specific pathogen recognition in *C. elegans*, worms mount highly specific transcriptional, immune, developmental, and behavioral responses upon exposure to different pathogenic species (Estes et al., 2010; McEwan et al., 2012; Moore et al., 2019; Wong et al., 2007; Zhang et al., 2005; Palominos et al., 2017). By examining how pathogens induce transgenerational learned avoidance in mothers and progeny, we sought to uncover how distinct pathogens are detected by hosts using non-canonical recognition pathways.

Here we have found that *C. elegans* can interpret the pathogenic status and bacterial species information through its ingestion and processing of bacterial small RNAs. This process utilizes both the canonical RNA interference and the piRNA pathways to convey information through the germline to neurons, inducing avoidance behavior in both mothers and in subsequent generations. We propose that this small RNA-based identification mechanism allows the animals not only to quickly adjust to pathogenic conditions, but also to predict future conditions that their progeny may experience.

## Results

### Small RNAs confer information about PA14 to *C. elegans*

To determine how *C. elegans* “identify” bacteria in a manner that is then utilized in the transgenerational inheritance of pathogenic avoidance, we tested bacterial components that the worms are exposed to during pathogenic training. Although bacterial metabolites can serve as cues that alter worm behavior (Meisel et al., 2014), we found that training worms with PA14 culture supernatant for 24 h did not induce avoidance learning (Fig 1A).

**Figure 1:**
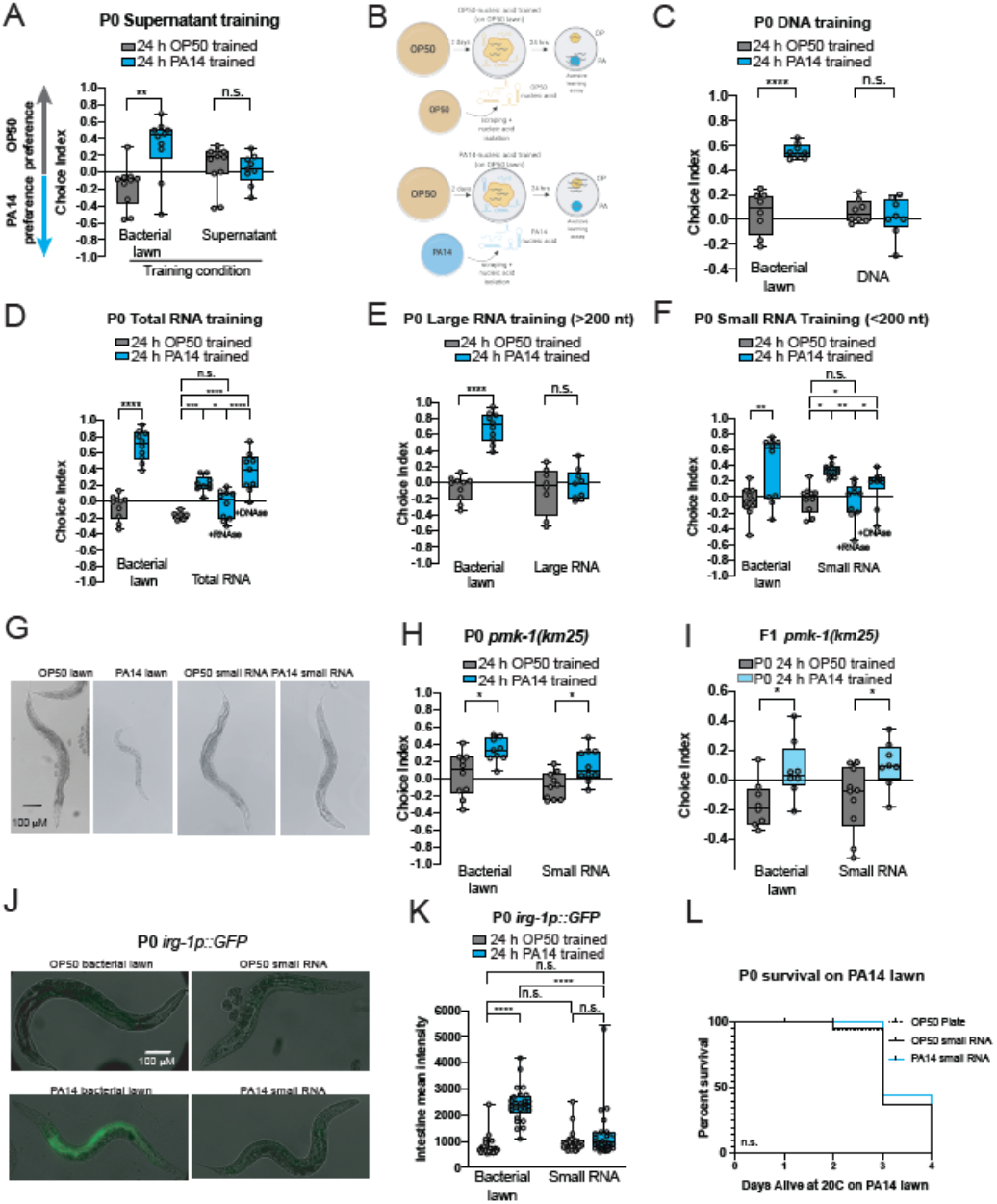
*Pseudomonas aeruginosa* (PA14) small RNAs are sufficient to induce *C. elegans* pathogen avoidance. (A) 24 h of training on supernatant from an overnight culture of PA14 is not sufficient to elicit maternal avoidance (P0) of PA14. 3 biological replicates were performed. (B) Small RNA training protocol and aversive learning assay for adult *C. elegans*. Choice index = (# of worms on OP50 - # of worms on PA14)/(total # of worms). (C) Total DNA purified from a PA14 bacterial lawns is not sufficient to elicit maternal avoidance (P0) of PA14. 2 biological replicates were performed. (D) Total RNA from PA14 bacterial lawns confers maternal avoidance of PA14. Total RNA treatment with RNAse abolishes this effect, while treatment with DNAse does not. 3 biological replicates were performed. (E) PA14 Large RNA (≥200 nt) training does not result in parental avoidance of PA14. 3 biological replicates were formed. (F) Small RNA (<200 nt) purified from PA14 bacterial lawns is sufficient to induce maternal avoidance of PA14. Small RNA treatment with RNAse abolishes this effect, while treatment with DNAse does not. At least 3 biological replicates were performed. (G) *C. elegans* exposed to PA14 small RNA are healthy compared to those exposed to PA14 bacterial lawns. (H) *pmk-1(km25)* is not required for PA14 bacterial lawn-induced or PA14 small RNA-induced pathogenic learning or (I) inherited behavior in the F1 generations. 2 biological replicates were performed. (I) *Irg-1p::GFP* expression is induced by PA14 bacterial lawn exposure, but not by PA14 small RNAs alone. (K) GFP intensity from (J) was quantified, n ≥ 26 worms per group. (F) PA14 small RNA trained P0s do not have a survival advantage on a lawn of PA14 compared to OP50 small RNA trained P0s. 1 biological replicate was performed. Aversive learning assays: For comparisons between training conditions of one genotype: One-Way ANOVA, Tukey’s multiple comparison test, mean ± SEM. n ≥ 6-10 choice assay plates with 50-200 worms per plate. For comparisons between multiple training conditions and genotypes: Two-Way ANOVA, Tukey’s multiple comparison test, mean ± SEM. n ≥ 6-10 choice assay plates with 50-200 worms per plate, *p ≤ 0.05, **p ≤ 0.01, ***p ≤ 0.001, ****p ≤ 0.0001, ns = not significant. Survival assay: Log-rank (Mantel-Cox) test, n = 80 worms per condition

We next hypothesized that worms might identify pathogens, such as PA14, instead through nucleic acid-encoded molecular codes. We isolated nucleic acid components from pathogenic (25°C, plate-grown) PA14, then tested them by adding the purified samples to spots of *E. coli* and exposing the worms for 24 h, followed by our standard choice assay (OP50 *E. coli* vs PA14; Fig. 1B) to determine which component might mimic pathogenic bacterial lawn exposure. These samples included total DNA (Fig. 1C), total RNA (Fig. 1D), large RNA (≥200nt) (Fig. 1E), small RNA (<200nt), and RNAse- and DNAse-treated fractions (Fig. 1D, F). The total RNA purified from plate-grown PA14, as well as DNAse-treated RNA, and small RNA samples, but not DNA or large RNAs, induced subsequent avoidance of PA14 in mothers (Fig. 1C-F, Supplement 1A-C). These results suggested that small RNAs are at least partially responsible for learned pathogenic avoidance. However, the avoidance induced by small RNA does not require virulence per se, as treatment of the worms with purified PA14 small RNA does not make them ill, unlike 24 h treatment with PA14 bacterial lawns (Fig. 1G). Moreover, the innate immune pathway regulator *pmk-1* (Kim, 2002; Troemel et al., 2006) is not required for the avoidance effect in mothers or progeny (Fig. 1H, I), nor are characteristic *pmk-1-* independent innate immune responses (*irg-1p::gfp*) induced by bacterial lawns of PA14 (Estes et al., 2010) (Fig. 1J, K). Exposure to small RNAs also does not improve survival on inescapable PA14 lawns (Fig. 1L). Therefore, small RNAs isolated from pathogenic PA14 are able to replicate the parental avoidance behavior normally induced by treatment with bacterial lawns, without exposure to the bacteria themselves, illness, or induction of the innate immune response.

Remarkably, we found that exposure of *C. elegans* to purified PA14 small RNA was sufficient to induce avoidance of PA14 not only in mothers, but also in the subsequent four generations (Fig. 2A, B; Supplemental Fig 2), completely mimicking the transgenerationally inherited learned avoidance effect of training on pathogenic bacteria lawns that we had previously identified (Moore et al., 2019). This avoidance effect persisted despite the fact that neither the mothers nor the progeny had ever encountered PA14. Together, these data suggest that the mechanism by which *C. elegans* transmits its learned avoidance does not depend on an innate immune response, or molecules canonically sensed by PRRs (e.g., proteins, lipopolysacchdarides, etc.), but rather on *C. elegans’* ability to identify the specific pathogen via bacterially-expressed small RNAs. Moreover, the similar magnitude of the behavior in progeny of live bacteria- and small RNA-trained animals suggests that the transgenerational response is due entirely to the PA14 small RNA response in worms (Fig. 2C).

**Figure 2:**
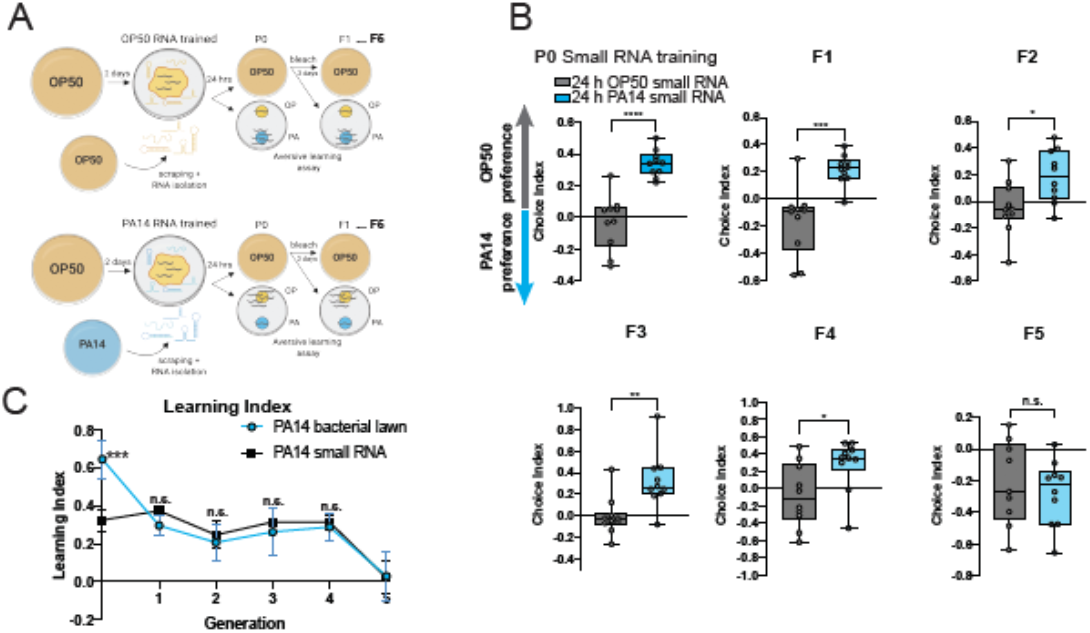
PA14 small RNAs induce transgenerational pathogen avoidance. (A) Small RNA training protocol for transgenerational inheritance of learned avoidance behavior. (B) P0 mothers exposed to PA14 small RNAs learn to avoid PA14. Untrained (naïve) progeny of PA14-small RNA trained mothers continue to avoid PA14 from generation F1-F4. 5^th^ generation progeny return to a state of PA14 attraction. Students t-test, mean ± SEM. n ≥ 6-10 choice assay plates with 50-200 worms per plate. At least two biological replicates were performed. (C) Learning index of bacterial lawn trained progeny and small RNA trained progeny at each generation. (Learning Index = Average OP50 choice index - average PA14 choice index). One-Way ANOVA, Tukey’s multiple comparison test, mean ± SEM. Statistics shown represent pairwise comparisons between PA14 bacterial lawn-trained mothers and PA14 small RNA-trained mothers for each generation, n ≥ 20 choice assay plates from two biological replicates with 50-200 worms per plate, *p ≤ 0.05, **p ≤ 0.01, ***p ≤ 0.001, ****p ≤ 0.0001, ns = not significant.

### Small RNAs are used by *C. elegans* to convey pathogenicity

*C. elegans* utilizes the information conveyed by the small RNA code of pathogenic PA14 to induce an avoidance response that starts in the mother and continues for four generations, despite never having been sick. To narrow down the set of small RNAs that might convey this information, first, we tested the maternal avoidance-inducing ability of small RNAs isolated from less-pathogenic growth conditions that do not induce transgenerational inheritance of pathogen avoidance, i.e., 15°C PA14 grown on plates (Moore et al., 2019), and from PA14 grown under planktonic (non-biofilm) conditions, i.e., overnight cultures of liquid-grown PA14 (Fig. 3A). Learned avoidance was specific for small RNAs isolated from bacterial lawns of 25°C-plate grown PA14 (Fig. 1F), as small RNAs isolated from less virulent 15°C-grown PA14 and liquid-grown PA14 did not induce subsequent avoidance of PA14 (Fig. 3B, C).

**Figure 3:**
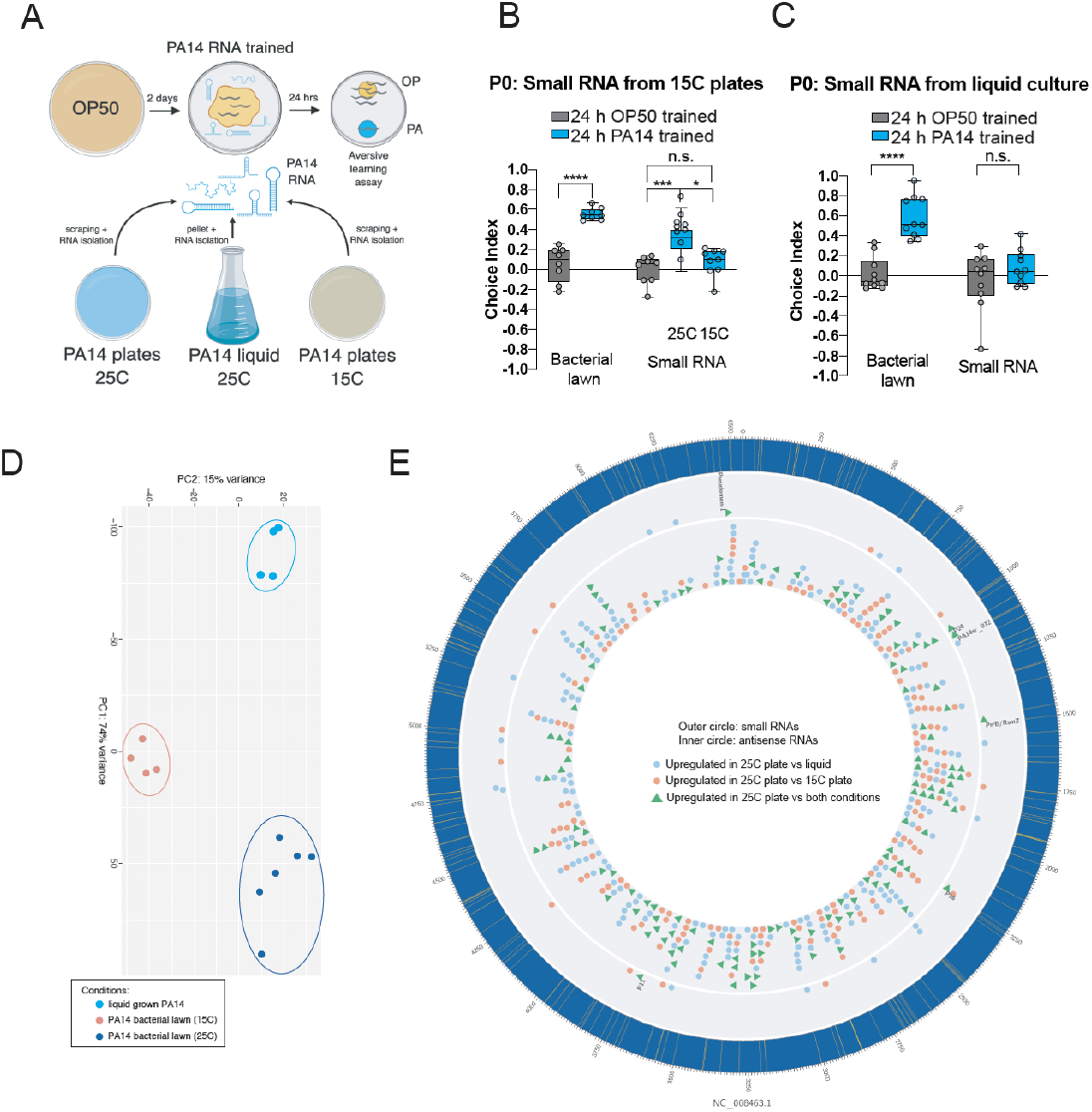
Small RNAs from PA14 are used by *C. elegans* to convey pathogenicity. (A) Small RNA training protocol using RNA isolated from PA14 cultures grown at 25°C or 15°C on plates, or in liquid culture. (B) Small RNAs isolated from less virulent PA14 (15°C) or (C) overnight liquid culture of PA14 do not cause maternal avoidance.One-Way ANOVA, Tukey’s multiple comparison test, mean ± SEM. n ≥ 6-10 choice assay plates with 50-200 worms per plate. Two-Way ANOVA, Tukey’s multiple comparison test, mean ± SEM. n ≥ 6-10 choice assay plates with 50-200 worms per plate, *p ≤ 0.05, **p ≤ 0.01, ***p ≤ 0.001, ****p ≤ 0.0001, ns = not significant. (D) PCA of small RNAs sequenced from PA14 under pathogenic (25°C) and less pathogenic conditions (liquid-grown and 15°C). (E) Differentially-expressed small RNAs and antisense RNAs (asRNA) in PA14 under pathogenic vs. less-pathogenic conditions. The outermost blue circle represents the PA14 genome, with previously identified (Wurtzel et al., 2012) small RNAs highlighted in yellow. Shown are small RNAs (outer circle) and asRNAs (inner circle) that were upregulated (DESeq2, adjusted p value ≤ 0.05) in the 25°C plate condition relative to 15°C plate grown PA14 (red) or liquid grown PA14 (blue). Overlapping sRNAs and asRNAs upregulated in the 25°C plate condition relative to both other conditions are shown as green triangles.

To identify the set of small RNAs that might convey the identity of pathogenic bacteria, we sequenced the small RNAs (including canonical small RNAs, possible novel small RNAs, and antisense RNAs (asRNA)) isolated from PA14 under the different growth conditions and identified those that were differentially expressed (Fig 3D-E). Of the previously annotated PA14 small RNAs (Wurtzel et al., 2012), 18 and 22 are significantly more highly expressed in 25°C-grown PA14 compared to 15°C-grown and liquid-grown PA14, respectively (Supplementary Tables 1, 2). Of these, six small RNAs are upregulated in the 25°C-grown samples compared to both conditions, including P11, P16, P24, PA14sr-032, Pseudomon, and PrrB/RsmZ (Fig. 3E, F, Supplementary Tables 1, 2). Several of these small RNAs, particularly Rsm genes, have been previously associated with or required for *Pseudomonas* virulence (Brencic and Lory, 2009; González et al., 2008; Heurlier et al., 2004), suggesting they may function as biomarkers of pathogenesis. We also found that 267 and 323 antisense RNAs are upregulated in 25°C-grown PA14 compared to 15°C-grown and liquid-grown PA14, respectively, with 120 asRNAs overlapping in both categories (Fig. 3E; Supplementary Tables 3, 4). In an additional unbiased approach to identify differences in small RNAs expressed in 25°C-grown PA14, sense and antisense RNA reads were compared across the entire ~6 Mb PA14 genome over every 100 bp window. Among the ~6,000-7000 upregulated changes observed in each 25°C-growth condition comparison, 1,705 regions were shared (Supplementary Tables 5, 6). Several of the differentially expressed regions correspond to the small RNA and asRNA changes identified above (Fig. 3E), but others correspond to mRNA changes, which also could act as post-transcriptional regulatory elements (Ren et al., 2017). These findings show that there is a set of small RNAs more highly expressed in the 25°C-plate growth conditions that could provide the worm with information to identify pathogenic bacteria and subsequently induce transgenerational pathogenic learning, essentially functioning as a signature of pathogenicity to the worms.

### The ASI neuron expresses *daf-7* in response to PA14 small RNAs

The DAF-7 TGF-β ligand is induced in the ASI and ASJ neurons upon direct exposure to PA14 (Meisel et al., 2014), and we previously found that *daf-7* is expressed at higher levels in the ASI neuron of F1 progeny of PA14-trained mothers (Moore et al., 2019). Upon exposure to PA14 small RNAs, *daf-7* is induced solely in the ASI neurons, resembling the expression pattern of progeny of trained mothers, despite being in the parental generation (Fig. 4A, B). Consistent with the role of PA14 small RNAs in TEI behavior, *daf-7* expression was elevated in F1 progeny of small RNA exposed mothers, as well (Fig 4. C, D).

**Figure 4:**
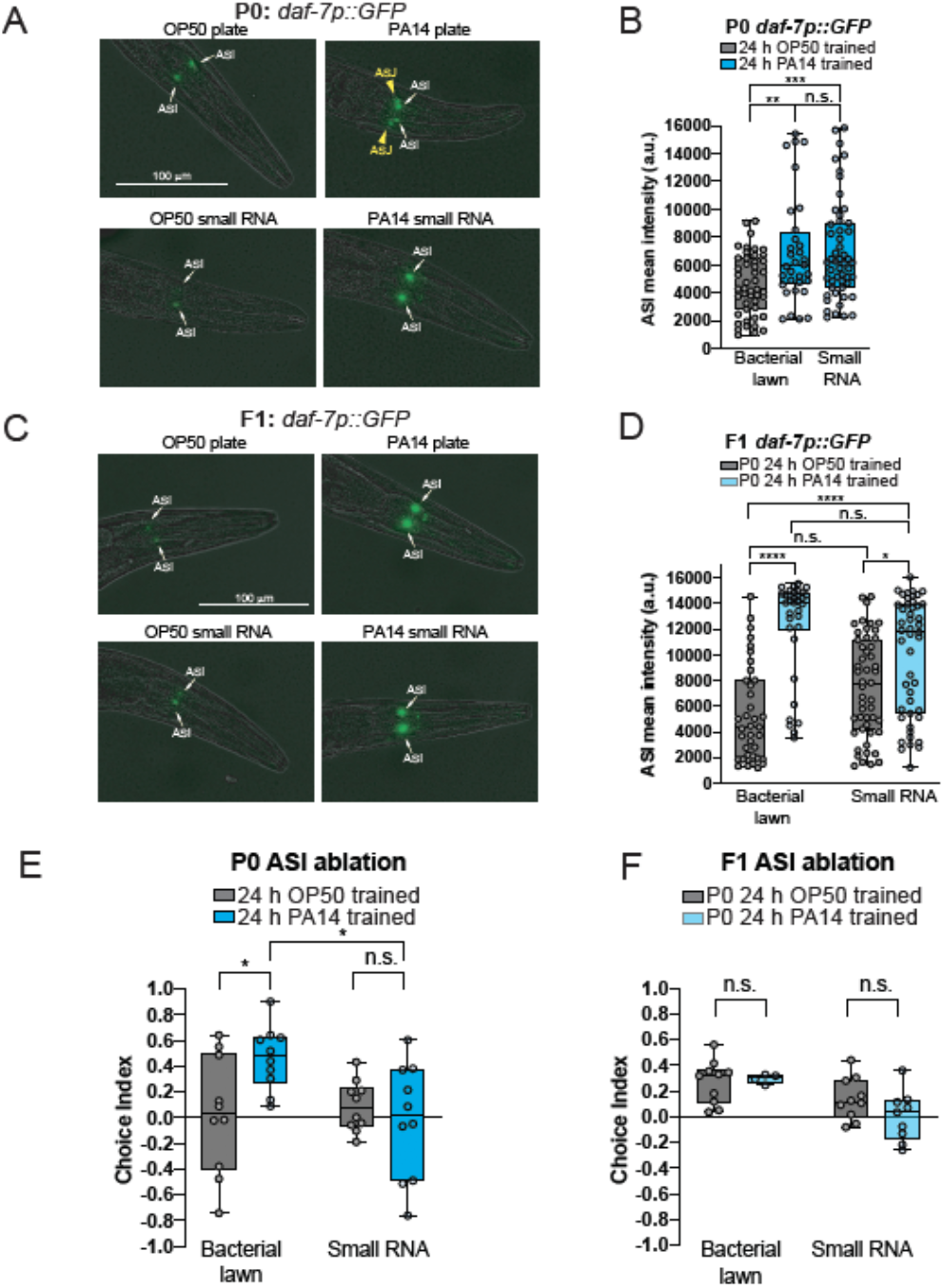
PA14 small RNAs are required for the parental and transgenerational neuronal changes in *daf-7*/TGF-β. **(A-B)** *daf-7::gfp* is expressed in the ASI of naïve animals (white arrows), and both the ASI and ASJ (yellow arrowheads) in PA14 plate-trained animals. PA14 small RNA exposure specifically increases *daf-7::gfp* in the ASI neurons. At least one biological replicate was performed. (C-D) *daf-7::gfp* expression remains elevated in both progeny (F1) of PA14 bacterial lawn trained mothers, and PA14 small RNA trained mothers. (E-F) The ASI neuron is required for parental (E) and transgenerational (F) small RNA induced pathogenic learning. At least one biological replicate was performed.One-Way ANOVA, Tukey’s multiple comparison test, mean ± SEM. n ≥ 6-10 choice assay plates with 50-200 worms per plate. For comparisons between multiple training conditions and genotypes: Two-Way ANOVA, Tukey’s multiple comparison test, mean ± SEM. n ≥ 6-10 choice assay plates with 50-200 worms, *p ≤ 0.05, ****p ≤ 0.0001, ns = not significant.

To assess the role of the ASI neuron in this process, we tested worms whose ASI neurons have been genetically ablated. While mothers trained on a bacterial lawn can learn to avoid PA14, likely through ASJ function as we previously showed (Moore et al., 2019), exposure of ASI-ablated worms to PA14 small RNAs does not induce avoidance of PA14 in trained mothers (Fig. 4E), or in their progeny (Fig. 4F). This suggests that the ASI neuron specifically responds to PA14 small RNAs to mediate transgenerational avoidance.

### The canonical RNA interference pathway is required for small RNA-induced avoidance

In RNA interference (RNAi), double-stranded RNA artificially expressed in bacteria is taken up by *C. elegans* and processed, and those small RNAs subsequently inhibit gene expression in the worm (Fire et al., 1998). We wondered whether the components of the canonical RNAi pathway (Ghildiyal and Zamore, 2009) are required for pathogenic avoidance induced by PA14 small RNAs. Indeed, while none of these genes are required for avoidance induced by direct bacterial lawn exposure to pathogen, we found that small RNA-induced avoidance requires the SID-1 (Shih and Hunter, 2011) and SID-2 (Braukmann et al., 2019; Winston et al., 2007) dsRNA transporters (Fig. 5A, B), as well as the RNAse III nuclease Dicer/DCR-1 that cleaves dsRNA for microRNA, primary exo-siRNA, and primary endo-siRNA biogenesis (Bernstein et al., 2001; Han et al., 2009; Ketting, 2001) (Fig. 5C, Supplement 4). Similarly, small RNA-induced avoidance requires the primary siRNA Argonaute RDE-1 (Tabara et al., 1999), RDE-2/MUT-8, a member of the MUTator complex that is essential for germline 22G siRNA biogenesis (Phillips et al., 2012; Tops, 2005), and RDE-4, a member of the ERI complex that binds long dsRNA to mediate exo-RNAi in the soma and germline by binding directly to DCR-1 (Parker, 2006; Tabara et al., 2002) (Fig. 5D-F). Together, these results suggest that the canonical RNA interference pathway is critical for the avoidance response induced by PA14 small RNAs.

**Figure 5:**
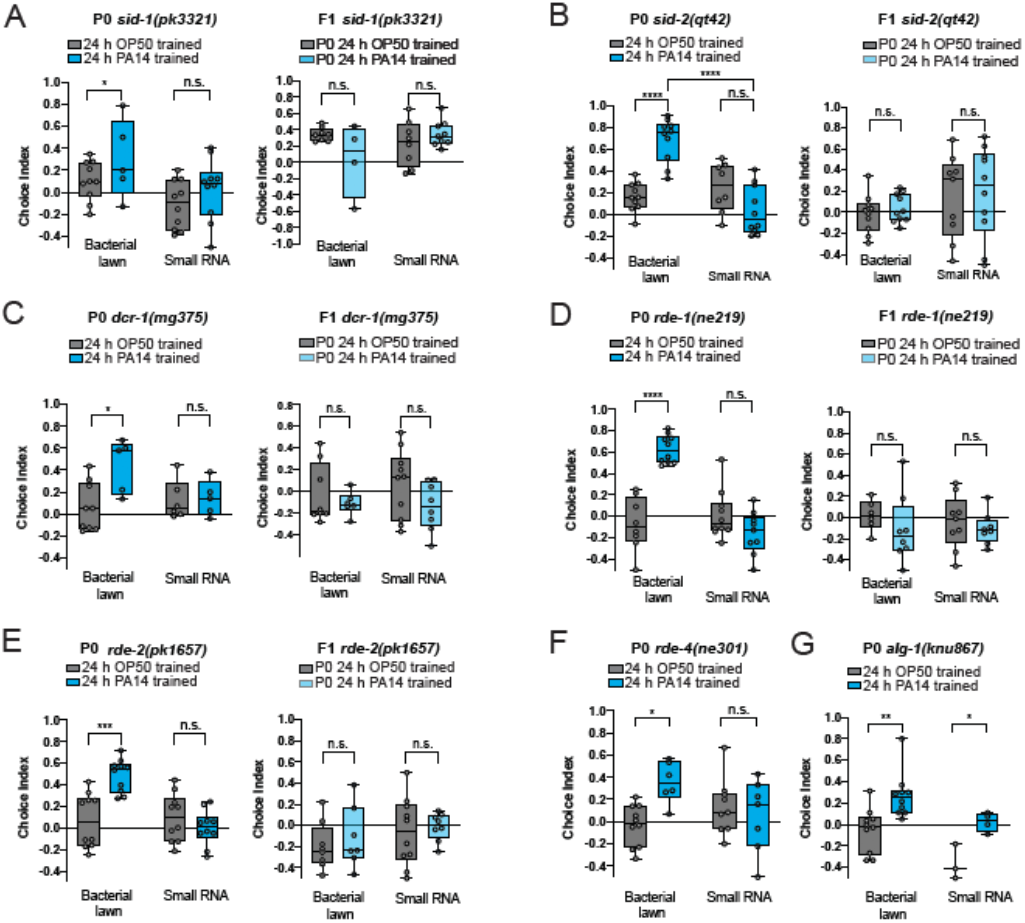
Worm dsRNA transport and processing machinery are required for bacterial sRNA-induced pathogen avoidance. (A-G) *sid-1(pk3321)* (A), *sid-2(qt42)* (B), *dcr-1(mg375)* (C), *rde-1(ne219)* (D), *rde-2(pk1657)* (E), *rde-4(ne301)*, and *alg-1(knu867)* mutants are able to learn to avoid PA14 after bacterial lawn training, but are specifically deficient in PA14 small RNA-induced learned avoidance. Progeny of *sid-1* (A), *sid-2* (B), *dcr-1* (C), *rde-1* (D), and *rde-2* (E) are defective in both transgenerational pathogen avoidance following maternal bacterial lawn and PA14 small training. One-Way ANOVA, Tukey’s multiple comparison test, mean ± SEM. n ≥ 6-10 choice assay plates with 50-200 worms per plate. For comparisons between multiple training conditions and genotypes: Two-Way ANOVA, Tukey’s multiple comparison test, mean ± SEM. n ≥ 6-10 choice assay plates with 50-200 worms. At least two biological replicates were performed, *p ≤ 0.05, **p ≤ 0.01, ***p ≤ 0.001, ****p ≤ 0.0001, ns = not significant.

By contrast, mutants of the microRNA-specific Argonaute *alg-1*/AGO-1 (Grishok et al., 2001) were able to learn avoidance upon exposure to both PA14 bacterial lawns and isolated small RNAs (Fig. 5G). Moreover, microRNA processing is unaffected in the *dcr-1(mg375)* mutant (Welker et al., 2010), which is unresponsive to small RNA from PA14 (Fig. 5C). Together, these data suggest that the secondary siRNA pathway, rather than the microRNA pathway, mediates PA14 small RNA-induced avoidance learning.

### The Piwi Argonaute piRNA pathway and a functional germline are required in mothers for small RNA-induced avoidance

We previously found that the piRNA regulator Piwi/PRG-1 Argonaute (Batista et al., 2008; Shirayama et al., 2012), its downstream RNAseD and RNA-dependent RNA polymerase components MUT-7 (Ketting et al., 1999) and RRF-1 (Aoki et al., 2007), respectively, as well as the SET-25 histone methylase (Towbin et al., 2012) and the HPL-2 chromatin reader (Couteau et al., 2002), are all required for the inheritance of learned pathogenic avoidance (Moore et al., 2019). However, we also previously showed that these proteins were not required for pathogen avoidance learning in mothers when trained on a bacterial lawn (Moore et al., 2019). Therefore, we were surprised to find that *prg-1* mutants were defective in the avoidance response after training on PA14 small RNA not only in F1 progeny, but in mothers themselves (Fig. 6A-B). Furthermore, *mut-7, rrf-1, rrf-3* (Pereira et al., 2019), and *hpl-2* mothers (P0) and progeny (F1) of PA14 small RNA-trained mothers had the same phenotype as *prg-1* mutants (Fig. 6C-F). These data suggest that the piRNA pathway is required not only for transgenerational inheritance of learned avoidance after bacterial lawn exposure, but also for the fraction of avoidance response mediated by small RNAs in mothers (Fig. 2C). Consistent with this hypothesis, *prg-1* mutant mothers exposed to PA14 bacterial lawns induced *daf-7* expression in the ASJ neurons (Fig. 6G; Supplemental Figure 4A), but failed to upregulate *daf-7* in the ASI neurons (Fig. 6G), which we have shown to be required for both transgenerational inheritance of avoidance (Moore et al., 2019), as well as small RNA-induced avoidance (Fig. 4). These data may explain why PA14 lawn-trained *prg-1* mothers can still learn to avoid PA14, but do not pass on this learned information, and cannot learn when exposed to PA14 small RNAs alone. That is, about half of the learned avoidance that mothers display may be due to innate immunity-like responses to PA14 (Meisel et al., 2014; Troemel et al., 2006; Zhang et al., 2005), but the remaining half of this response is due to small RNA-induced avoidance, and this latter process is utilized in transgenerational inheritance of learned avoidance.

**Figure 6:**
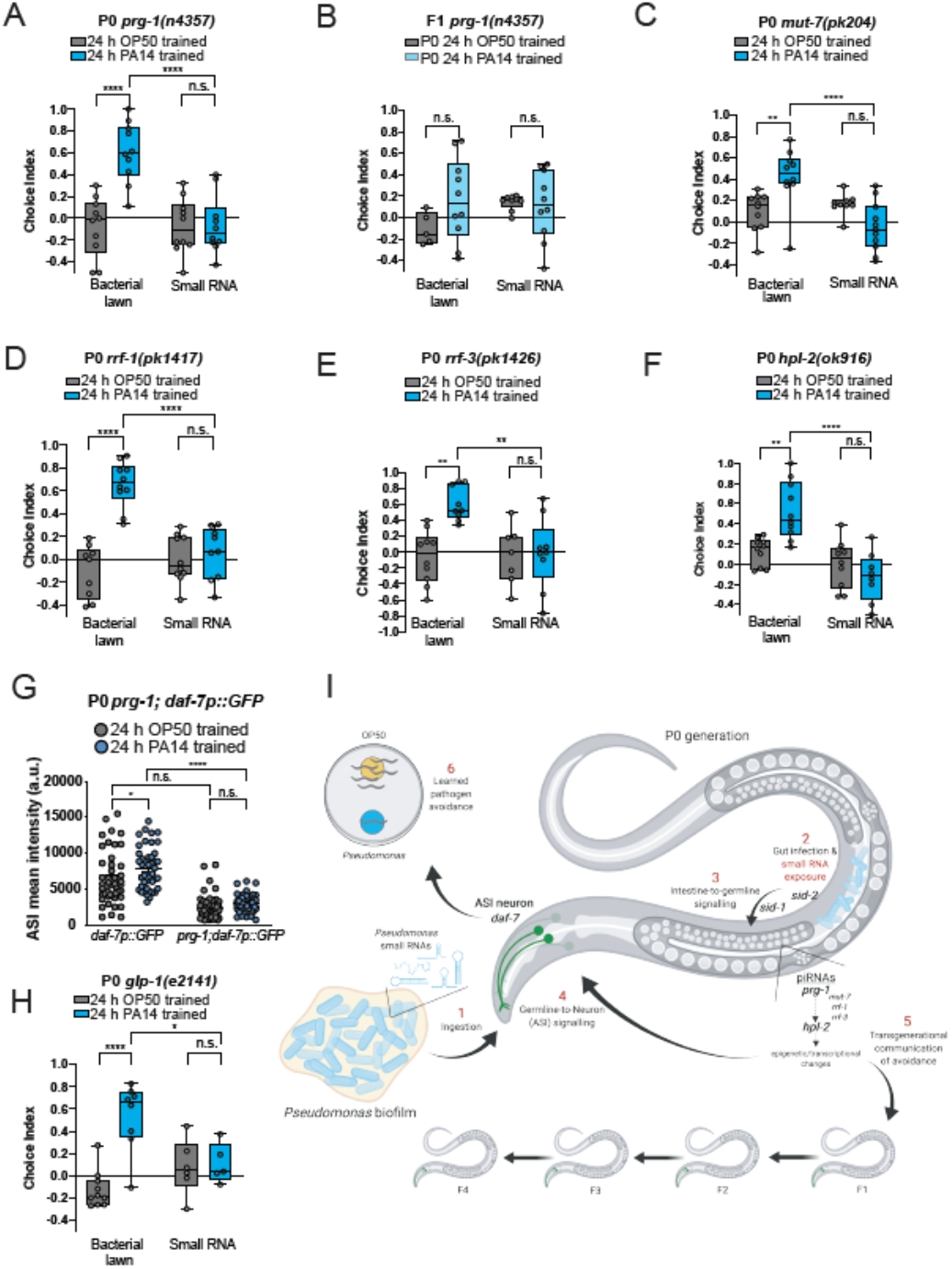
The *C. elegans* germline and PRG-1/PIWI and 22G-RNA pathways are required for sRNA-induced pathogen avoidance. **(A)** *prg-1(n4357)* mutants can learn to avoid PA14 after training on a bacterial lawn, but are defective in pathogenic learning when trained on PA14 small RNAs. (B) *prg*-1 is required for inheritance of both PA14 bacterial lawn-training and PA14 small RNA-training induced aversive learning. (C-F) *mut-7(pk204)* (C), *rrf-1(pkl417)* (D), *rrf-3(pkî426)* (E), and *hpl-2(ok916)* (F) are not required for bacterial lawn aversive learning, but are specifically required for PA14 small RNA-mediated learned avoidance in P0 mothers. (G) *prg-1* mutants, like wild-type worms (Fig. 4), induce *daf-7::gfp* expression in the ASJ neuron, but not the ASI neuron, after PA14 bacterial lawn exposure. (<u>H</u>)*glp-1(e2141)* mutants lacking a germline (i.e., raised at 25°C) have normal naïve preference and can learn to avoid PA14 after training on a bacterial lawn, but are defective in small RNA-induced pathogenic learning. Three biological replicates were performed. One-Way ANOVA, Tukey’s multiple comparison test, mean ± SEM. n ≥ 6-10 choice assay plates with 50-200 worms per plate. For comparisons between multiple training conditions and genotypes: Two-Way ANOVA, Tukey’s multiple comparison test, mean ± SEM. n ≥ 6-10 choice assay plates with 50-200 worms. At least one biological replicate was performed, *p ≤ 0.05, **p ≤ 0.01, ***p ≤ 0.001, ****p ≤ 0.0001, ns = not significant. (J) Model of PA14 small RNA-induced transgenerational learned avoidance.

### A functional germline is required for small RNA-induced avoidance

Because PRG-1 is required for *daf-7* expression changes in the ASI (Fig. 6G), we next asked whether bacterial small RNAs act directly in neurons, as proposed in recent work (Posner et al., 2019), or whether signaling through other tissues is required to induce gene expression in the nervous system and subsequent behavioral changes. Therefore, we examined the requirement of the germline in PA14 small RNA-mediated avoidance learning using *glp-1(2141)* mutants, which lack a germline when grown at the restrictive temperature (25°C) (Austin and Kimble, 1987). We found that animals lacking a germline can learn to avoid PA14 in a choice assay after training on a PA14 bacterial lawn (Fig. 6H, left), demonstrating that innate immunity learning mechanisms do not rely on a functional germline. However, *glp-1* mutants fail to exhibit small RNA-induced avoidance of PA14 (Fig. 6H, right), suggesting that activity in the germline is required specifically for small RNA induction of this neuronally-directed avoidance behavior. Thus, bacteria-derived small RNAs are likely not acting directly in neurons, but rather through an indirect mechanism that requires an intact germline to communicate to neurons.

### A model of small RNA-induced transgenerational pathogen avoidance

Our data suggest a model in which pathogen avoidance is induced by at least two pathways: worms respond acutely (within 4 h) through the innate immune response (Troemel et al., 2006) to pathogen exposure and metabolites (Meisel et al., 2014; Pereira et al., 2019; Zhang et al., 2005), which induces *daf-7* expression in the ASJ neuron, resulting in about half of the avoidance behavior exhibited by trained mothers (P0s; Fig. 2C). In a separate pathway that we have identified here, 24 h exposure to pathogenic bacteria and subsequent uptake of bacterial small RNAs in the intestine (Fig. 6I), followed by processing in the germline through the canonical RNA interference pathway and piRNA processing (Batista et al., 2008) to HPL-2, induces *daf-7* expression in the ASI neuron, ultimately contributing not only avoidance of PA14, but also to its progeny (Fig. 2C; 6I). Despite the brevity of the exposure, this latter pathway is propagated for four additional generations and maintains the same magnitude of avoidance until F5 (Fig. 2C).

There appears to be a correlation between the transgenerational effect and the response to small RNAs; that is, all the genes that we found to be required for the small RNA-induced response in mothers (P0) are also required for the transgenerational inheritance of learned pathogen avoidance. Therefore, we propose that the set of small RNAs from PA14 cultivated under pathogenic conditions is sufficient to engage a pathway that accounts for half of the mother’s avoidance response, and for the entire transgenerationally-inherited avoidance response (F1-F4)(Fig. 2C; 6I).

### Small RNA-induced pathogen avoidance is species-specific

Previously, Zhang et al., 2005 showed that exposure to *Serratia marcescans* induces pathogenic avoidance, but we found that this effect is limited to the parental generation, suggesting that inheritance of learned pathogenic avoidance is not induced by all pathogens (Moore et al., 2019). Although training on pathogenic *S. marcesans* lawns induces avoidance, small RNAs isolated from *S. marcesans* do not induce avoidance (Fig. 7A), consistent with our observed correlation between transgenerational inheritance of avoidance behavior and maternal avoidance response to small RNAs.

**Figure 7:**
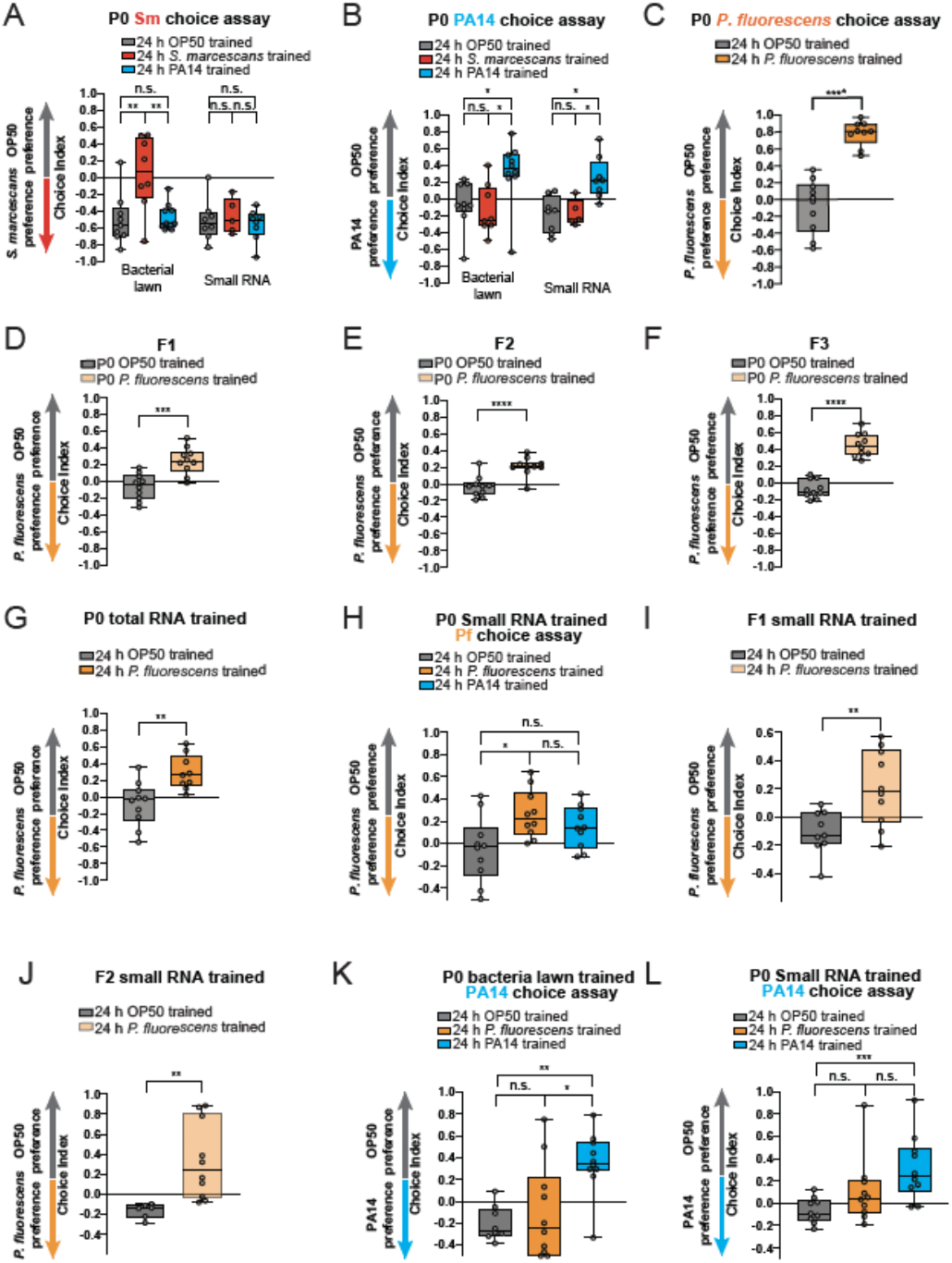
*C. elegans* uses small RNAs to transgenerationally avoid environmentally abundant and pathogenic bacteria. (A) *C. elegans* can specifically learn to avoid *S. marcescans* (Sm) after training on Sm (red) compared to training on PA14 (blue). Small RNAs isolated from Sm or PA14 are not sufficient to to elicit avoidance of Sm. (B) *C. elegans* specifically learn to avoid PA14 after training on either PA14 bacterial lawns or PA14 small RNAs. One-Way ANOVA, Tukey’s multiple comparison test, mean ± SEM. n ≥ 6-10 choice assay plates with 50-200 worms per plate. For comparisons between multiple training conditions and genotypes: Two-Way ANOVA, Tukey’s multiple comparison test, mean ± SEM. n ≥ 6-10 choice assay plates with 50-200 worms. At least two biological replicates were performed. (C) 24 h of training on *P. fluorescens* (Pf) bacterial lawns is sufficient for maternal learned avoidance of Pf. (D-F) Untrained (naive) progeny of Pf-trained mothers avoid Pf in generations FI (D), F2 (E), and F3 (F). (G) Exposure to *P. fluorescens* total RNA is sufficient to induce maternal avoidance of Pf, similar to Pf bacterial lawn training. Students t-test, mean ± SEM. n ≥ 6-10 choice assay plates with 50-200 worms per plate. At least 2 biological replicates were performed. (H) Pf Small RNAs (<200nt) are sufficient to elicit P0 avoidance of Pf (orange), while PA14 small RNA training (blue) does not change the worms’ preference to Pf. One-Way ANOVA, Tukey’s multiple comparison test, mean± SEM. n ≥ 6-10 choice assay plates with 50-200 worms per plate. For comparisons between multiple training conditions and genotypes: Two-Way ANOVA, Tukey’s multiple comparison test, mean ± SEM. n ≥ 6-10 choice assay plates with 50-200 worms. At least two biological replicates were performed. (I-J) Progeny (FI, I) and grandprogeny (F2, J) of Pf small RNA trained mothers avoid Pf. (K) 24 h of training on a PA14 bacterial lawn (blue) induces learned avoidance of PA14, while training on a Pf bacterial lawn (orange) does not change the worms’ preference to PA14. Students t-test, mean± SEM. n ≥ 6-10 choice assay plates with 50-200 worms per plate. At least 2 biological replicates were performed.(G) 24 h of training on PA14 small RNAs (blue), but not Pf small RNAs (orange), confers PA14 learned avoidance. One-Way ANOVA, Tukey’s multiple comparison test, mean ± SEM. n ≥ 6-10 choice assay plates with 50-200 worms per plate. For comparisons between multiple training conditions and genotypes: Two-Way ANOVA, Tukey’s multiple comparison test, mean ± SEM. n ≥ 6-10 choice assay plates with 50-200 worms. At least two biological replicates were performed.

Why might *C. elegans* induce multi-generational avoidance of some pathogens, but not others? We surmised that worms may have evolved a mechanism to respond to pathogens that 1) have similar-smelling relatives that are beneficial, and 2) are abundant in *C. elegans’* environment. *S. marcescans* might not have induced such a mechanism because worms may not frequently encounter this pathogen or may have no relatives that are beneficial to *C. elegans* and “smell” the same to the worms. By contrast, *Pseudomonas* species make up perhaps a third of the *C. elegans* microbiome (Dirksen et al., 2016; Samuel et al., 2016), and at least a quarter of the bacterial species in their natural environment; thus, worms are likely to encounter both beneficial and detrimental *Pseudomonas* species quite frequently. We hypothesized that this small RNA mechanism should only function in species that are both abundant and potentially both beneficial and detrimental to *C. elegans*.

To test this hypothesis, we asked whether the behavioral changes associated with small RNA-induced pathogen avoidance were specific to PA14, or whether generalized changes occur that alter attraction to other known *C. elegans* pathogens. Similar to our previous results obtained by training on bacterial lawns (Moore et al., 2019), worms trained on PA14 small RNAs specifically avoided PA14 but not *S. marcescens* (Fig. 7B), suggesting that the learned avoidance is a response specific to the bacteria from which the small RNAs were obtained.

In contrast to *S. marcesans*, the *Pseudomonas* species *P. fluorescens* (Pf), which can be either beneficial or detrimental to *C. elegans* (Burlinson et al., 2013; Tan et al., 1999) is abundant in the set of *Pseudomonas* species present both in the worm’s environment and its microbiome (Dirksen et al., 2016; Kissoyan et al., 2019; Samuel et al., 2016). We found that *C. elegans* both learn to avoid *P. fluorescens* in the maternal generation (Fig. 7C) and continue to avoid *P. fluorescens* for several generations (F1-F3) (Fig. 7D-F, Supplemental Fig. 4A-D). To test the model that small RNAs are sufficient to induce transgenerational avoidance behavior, we isolated total and small RNAs from *P. fluorescens*, and tested their ability to induce avoidance in mothers and progeny. We found that training with either total (Fig. 7G) or small RNAs (Fig. 7H) from *P. fluorescens* caused mothers to avoid *P. fluorescens*, and this behavior persisted in the progeny for at least two generations (Fig. 7I-J, Supplement 4E-I). Training with *P. fluorescens* small RNAs induced avoidance that was specific to *P. fluorescens*, as Pf small RNA-trained worms exhibited attraction to PA14, rather than avoidance (Fig. 7H). Similarly, PA14 bacterial lawn (Fig. 7K) or small RNA-trained (Fig. 7L) worms learned to avoid PA14, but display attraction to Pf rather than avoidance. These results demonstrate that small RNAs from multiple *Pseudomonas* species can be detected by *C. elegans*, and that information is transmitted for several generations, resulting in bacterial species-specific behavioral responses.

## Discussion

### *C. elegans* can identify bacteria by “reading” their small RNAs

Cellular responses to exogenous RNA were discovered over 20 years ago in *C. elegans* (Fire et al., 1998), but the natural context in which worms evolved the ability to detect and respond to RNA from the environment remains unclear (Braukmann et al., 2019), especially since systemic and transgenerational RNAi pathways are not required for defense against Orsay virus, the only known natural viral pathogen of *C. elegans* (Ashe et al., 2015). Here we have identified a trans-kingdom signaling system that does not rely on secreted molecules, as has been previously reported (Meisel et al., 2014; Papenfort and Bassler, 2016), but rather on “reading” of a bacteria’s small RNAs, particularly the set that are required for surface-grown PA14 virulence (S. Chuang & Z. Gitai, pers. comm.). Previously, trans-kingdom signaling has been reported in which small RNAs of a pathogen hijack the host immune system to avoid detection (Li et al., 2014; Weiberg et al., 2013). By contrast, we have found that *C. elegans* uses bacterial small RNAs to mount a species-specific avoidance response that is propagated for four generations. This small RNA-sensing pathway depends on processing through the germline, and is independent of the pathways induced by innate immune systems and bacterial metabolites. The phenomenon described here may explain why both systemic and TEI responses to exogenous RNA exist: to modify the worm’s behavior in response to encounters with naturally abundant and pathogenic bacterial species, which are identified by the worm through unique small RNA signatures. While other previously-described transkingdom signaling systems are beneficial to the pathogen, *C. elegans* identifies small RNAs of pathogens in order to protect itself and to induce the search for less pathogenic food sources. The fact that this avoidance endures for several generations suggests that an acute response is not sufficient to protect against some abundant pathogens, but must be transient enough to be reset in order to prevent account for changes in available bacterial food quality.

### The physiological relevance of small RNA-induced pathogen avoidance

An emerging model of transgenerational inheritance of behavior suggests that small RNA signals originating from neurons are communicated to the germline and subsequently to progeny (O’Brien et al., 2019; Posner et al., 2019). Our data support a different model of transgenerational inheritance, in which primary detection of a small RNA signal in the intestine and subsequent activity in the germline communicates a signal to neurons to induce behavioral changes. Worm neurons are largely resistant to RNAi because they lack expression of *sid-1* and *sid-2* dsRNA transporters (Braukmann et al., 2019; Winston et al., 2002, 2007). It is therefore not surprising that, in the context of *Pseudomonas-small* RNA-mediated pathogen avoidance, neurons are not the tissue where primary detection of bacterial small RNA occurs. Whether other animals utilize their RNA interference pathways for similar purposes will be interesting to explore. Such a mechanism might be particularly useful in tissues that are in constant contact with bacteria, such as mammalian intestinal cells exposed to the microbiome.

The process we have uncovered here is also distinct from developmental responses to bacterial exposure, particularly prolonged, multi-generational exposure that induces dauer formation (Palominos et al., 2017; Moore and Murphy, unpublished data). When worms are forced to encounter pathogenic *Pseudomonas* continuously (≥ 6 days) over multiple generations, eventually their progeny halt development at the alternative, pre-reproductive L3 larval stage known as dauer, undergoing extensive remodeling of their cuticle, mouths, and other tissues. Dauer remodeling is extremely energy-intensive and reproductively unfavorable - a last-ditch effort to survive under extreme conditions - which may be why it requires such a prolonged exposure to pathogens to induce this extreme response. By contrast, 24h of exposure to *Pseudomonas* is sufficient to cause worms to switch their behavior from attraction to avoidance, and to induce the transgenerational inheritance of this learned avoidance. Rapid and lasting behavioral changes may be a more efficient and reproductively advantageous mechanism to force the worms to escape and thus explore a new and potentially safe environment. By passing this avoidance behavior on to several generations of their progeny, they may spare them from ever experiencing a prolonged exposure to the same pathogen, despite its potential abundance in the environment. Such a species-specific and plastic response may provide worms with a powerful survival mechanism that is fast-acting and potentially rapidly reversible (Houri-Ze’evi et al., 2019), a first line of defense against pathogens. It will be interesting to determine whether the dauer induction caused by prolonged pathogen exposure, which has been shown to also require some components of the RNAi machinery (Palominos et al., 2017) is an extension of the process that initially induces the rapid transgenerational avoidance effect we have described here, or whether it is an independent process.

### Bacterial small RNA-induction of transgenerational learned avoidance: a nascent adaptive immune system?

Bacterial small RNAs act as a decipherable code that allows the worms and their progeny to identify pathogens and respond appropriately when confronted with the pathogen in the future, prior to actually becoming ill. That is, the progeny already know to avoid these pathogens not because they have become sick, but because they have been “vaccinated” against the pathogen. This nascent adaptive immune system may enable *C. elegans* to identify and avoid pathogenic bacteria that exist in high abundance in their microbiome, and to pass on this avoidance to several generations of progeny. This trans-kingdom communication paradigm may represent a form of *C. elegans’* adaptive ‘immune’ memory that prepares future generations for encounters with harmful environmental conditions, allowing them to properly respond to a pathogenic threat.

## Methods

### C. elegans and bacterial strains cultivation

Worm strains were provided by the *C. elegans* Genetics Center (CGC). FK181: *ksls2 [Pdaf-7::GFP + rol-6(su1006)]*, CF1903: *glp-1(e2141)*, PY7505: oyls84 [gpa-4p::TU#813 + *gcy-27p*::TU#814 + *gcy-27p*::GFP + *unc-122p*::DsRed], NL3321: *sid-1(pk3321)*, HC271: ccls4251 !; qtls3 sid-2(qt42) III, mls11 IV, YY470: *dcr-1(mg375)*, SX922: *prg-1(n4357)*, WM27: *rde-1(ne219)*, NL3531: *rde-2(pk1657)*, WM49: *rde-4(ne301)*, MAH23: *rrf-1(p1417)*, NL2099: *rrf-3(ok1426)*, NL917: *mut-7(pk204)*, RB995: *hpl-2(ok916)*, COP2012: *alg-1(knu867)*, KU25: *pmk-1(km25)*, AU133: *agls17[Pmyo-2::mCherry + Pirg-1::GFP]*. CQ605 *prg-1(n4357) I; ksIs2 [Pdaf-7::GFP + rol-6(su1006)]*. OP50 was provided by the CGC. *S. marcescens* (ATCC 274) was purchased from ATCC. PA14 was a gift from Z. Gitai, and *P. fluorescens pf15* was a gift from M. Donia. Worm strains were maintained at 15°C on High Growth Media (HG) plates (3 g/L NaCl, 20 g/L Bacto-peptone, 30 g/L Bacto-agar in distilled water, with 4 mL/L cholesterol (5 mg/mL in ethanol), 1 mL/L 1M CaCl2, 1 mL/L 1M MgSO4, and 25 mL/L 1M potassium phosphate buffer (pH 6.0) added to molten agar after autoclaving) on *E. coli* OP50 using standard methods. Bacteria strains were cultured overnight in autoclaved and cooled Luria Broth (10 g/L typtone, 5 g/L yeast extract, 10 g/L NaCl in distilled water) shaking 37°C 250 rpm.

### Pathogen training

Eggs from young adult hermaphrodites were obtained by bleaching and placed onto High Growth (HG) plates and left at 20°C for 2 days. For bacterial lawn training, plates were prepared by inoculating overnight cultures of OP50 and pathogen in LB at 37°C. Overnight cultures were diluted in LB to an Optical Density (OD_600_) = 1 and used to fully cover Nematode Growth Media (NGM) plates. To prepare small RNA training plates 200 μL of OP50 was spotted in the center of a NGM. All plates were incubated at 25°C in separate incubators for 2 days. On day of training (i.e., 2 days post bleaching) plates were left to cool on a bench top for < 1 hr. Immediately before plating of worms the following was prepared; supernatant training: 1 mL of filtered supernatant was placed on top of OP50 spots and left to completely dry, DNA training: 10 ng of DNA was placed on top of OP50 spots, 10 μL of pooled L4 worms were plated onto OP50 fully seeded, while 5 μL of worms onto small RNA spotted training plates, and 40 μL of worms were plated onto pathogen seeded training plates. Worms were incubated on training plates at 20°C in separate containers for 24 h. After 24 h, worms were washed off plates in M9 3x. Some worms were used for an aversive learning assay, while the majority of worms were bleached onto HG plates at 20°C for 3 days.

### Aversive learning assay

Overnight bacterial cultures were diluted in LB to an Optical Density (OD600) = 1, and 25 μL of each bacterial suspension was seeded on a 60 mm NGM plate and incubated at 25°C for 2 days. After 2 days assay plates were left at room temperature for 1 h before use. Immediately before use, 1 μL of 1M sodium azide was spotted onto each bacterial lawn to be used as a paralyzing agent during choice assay. To start the assay (modified from (Zhang et al., 2005)), worms were washed off training plates in M9, and washed 2 additionally times in M9. 5 μL of worms were spotted at the bottom of the assay plate, using a wide orifice tip, midway between the bacterial lawns. Assays were incubated at room temperature for 1 h before counting the number of worms on each lawn.

In experiments in which F1 and subsequent generations are used: All animals tested are washed off HG plates with M9 at Day 1. Some of the pooled animals are subjected to an aversive learning assay, while the majority of worms are bleached to obtain eggs, which were then placed onto HG plates left at 20°C for 3 days and used to test F2s.

### Bacterial growth and training plate preparation

All bacterial samples were prepared by inoculating overnight cultures of pathogen in LB at 37C.

DNA: Overnight cultures were pelleted at 5,000 x g for 10 minutes at room temperature. DNA was prepared from pelleted bacteria according using the Qiagen DNeasy Blood and Tissue kit and subsequently used fresh.

Liquid: 1 mL of overnight cultures (undiluted) were put onto OP50 spots and left to completely dry at room temperature. Supernatant: Overnight cultures (undiluted) were pelleted at 5,000 x g for 10 minutes at room temperatures. Supernatant was removed and filtered using a 0.22 μm filter. 1 mL of filtered supernatant was put onto OP50 spots and left to completely dry at room temperature.

Total, large, and small RNA and DNAse/RNAse treated RNA: Bacterial lawns were collected from the surface of NGM plates using a cell scraper. Briefly, 1 mL of M9 buffer was applied to the surface of the bacterial lawn, and the bacterial suspension following scraping was transferred to a 15 mL conical tube. PA14 from 10 plates or OP50 from 15 plates was pooled in each tube and pelleted at 5,000 x g for 10 minutes at 4°C. The supernatant was discarded and the pellet was resuspended in 1 mL of Trizol LS for every 100 μL of bacterial pellet recovered. The pellet was resuspended by vortexing and subsequently frozen at −80°C until RNA isolation. 240 mg of Total RNA or RNAse/DNAse treated Total RNA, 100 mg of small or large or RNAse/DNAse treated small RNA was placed directly onto OP50 spots and left to completely dry at room temperature before use.

### Bacteria RNA isolation

To isolate RNA from bacterial cultures, Trizol lysates were incubated at 65°C for 10 min with occasional vortexing.

Debris was pelleted at 7,000 x g for 5 min at 4°C. The supernatant was transferred to new tubes containing 1/5 the volume of chloroform. Samples were mixed thoroughly by inverting and centrifuged at 14,000 x g for 10 min at 4°C. The aqueous phase was used at input for RNA purification using the mirVana miRNA isolation kit according to the manufacturer’s instructions for total RNA, large RNA (>200 nt), and small RNA (<200 nt) isolation. Purified RNA was frozen at −80°C until further use.

For RNase treatments of purified RNA, samples containing 100μg of RNA were treated with 2.5 μL of an RNAse A 500 U/mL and RNase T 20,000 U/mL cocktail for every 50 μL of RNA (RNase Cocktail Enzyme Mix, Ambion). Samples were incubated at room temperature for 20 min before adding to worm training plates seeded with OP50. RNase degradation was confirmed using an Agilent 2100 Bioanalyzer.

For DNAse treatment, samples containing 100μg of purified RNA were treated with 2U of DNAseI per 10 μg of RNA using the Invitrogen DNA-free kit according to the manufacturer’s instructions. DNAse-treated RNA samples were added to OP50-seeded plates as described above.

### Imaging and fluorescence quantitation

Z-stack multi-channel (DIC, GFP) of day 1 adult GFP transgenic worms were imaged every 1 μm at 60X magnification; Maximum Intensity Projections and 3D reconstructions of head neurons were built with Nikon *NIS-Elements*. To quantify *daf-7p::GFP* levels, worms were prepared and treated as described for pathogen training. Worms were mounted on agar pads and immobilized using 1 mM levamisole. GFP was imaged at 60X magnification and quantified using *NIS-Elements* software. Average pixel intensity was measured in each worm by drawing a bezier outline of the neuron cell body for 2 ASI head neurons and/or 2 ASJ head neurons.

### Small RNA sequencing

The size distribution of small RNA samples were examined on Bioanalyzer 2100 using RNA 6000 Pico chip (Agilent Technologies, CA). For small RNA-seq, around 300 nanogram of total RNA from each sample was first treated with RNA 5’ Pyrophosphohydrolase (New England Biolabs, MA) at 37°C for 30 minutes, then converted to Illumina sequencing library using the PrepX RNA-seq library preparation protocol on the automated Apollo 324™ NGS Library Prep System (Takara Bio, CA). Basically, the treated RNA samples were ligated to 2 different adapters at each end, then reverse transcribed to cDNA and amplified by PCR using different barcoded primers. The libraries were examined on Bioanalyzer DNA High Sensitivity chips (Agilent, CA) for size distribution, quantified by Qubit fluorometer (Invitrogen, CA), then pooled at equal molar amount and sequenced on Illumina NovaSeq 6000 S Prime flowcell as single-end 122 nt reads. The Pass-Filter (PF) reads were used for further analysis.

### Small RNA analysis

*Pseudomonas aeruginosa* (UCBPP-PA14) small RNA stranded reads were trimmed to remove adapters using Cutadapt (v1.16.6). Reads were mapped to the CP000438.1 genome using BWA-MEM. For small RNA analysis, count tables were generated using previously annotated intergenic non-coding RNAs (sRNA). Differential gene expression between 25 °C plate vs 15 °C plate and 25 °C plate vs liquid conditions was performed using DESeq2. For antisense RNA (asRNA) analysis, reads were mapped to the reverse strand of coding regions, and DESeq2 was performed as described above. To compare forward and reverse RNA abundance over every 100 bp genome interval, mean per-window depth was calculated using the mosdepth software (Pedersen and Quinlan, 2018) for forward and reverse mapping reads separately. The mean counts for each window were rounded to the nearest integer and imported into R 3.6.0 (R Core Team, 2019). The counts were transformed using the variance stabilizing transform (vst) function in DESeq2 for use in sample comparisons (heatmap and PCA). Differential expression was calculated using DESeq2 1.24.0 (Love et al., 2014) using default parameters. The lfcShrink function was used with the adaptive shrinkage estimator from the ashr’ package (Stephens, 2016) was used to shrink the log_2_ fold change estimates for windows with low counts to improve visualization.

## Acknowledgments

We thank the *C. elegans* Genetics Center for strains; the Genomics Core Facility at Princeton University, J. Wiggins, and J. Miller for RNA-seq library preparation and sequencing. We thank W. Wang for helping develop methods to sequence bacterial small RNAs; Z. Gitai and M. Donia for bacteria strains; and Z. Gitai and the Murphy lab for discussion. CTM is the Director of the Glenn Center for Aging Research at Princeton and an HHMI-Simons Faculty Scholar.

## Author contributions

RK, RSM, and CTM designed experiments. RSM and RK performed experiments and analyzed data. LP and RK analyzed small RNAseq data. RK, RSM, and CTM wrote the manuscript.

---

**Supplemental Figure 1:**
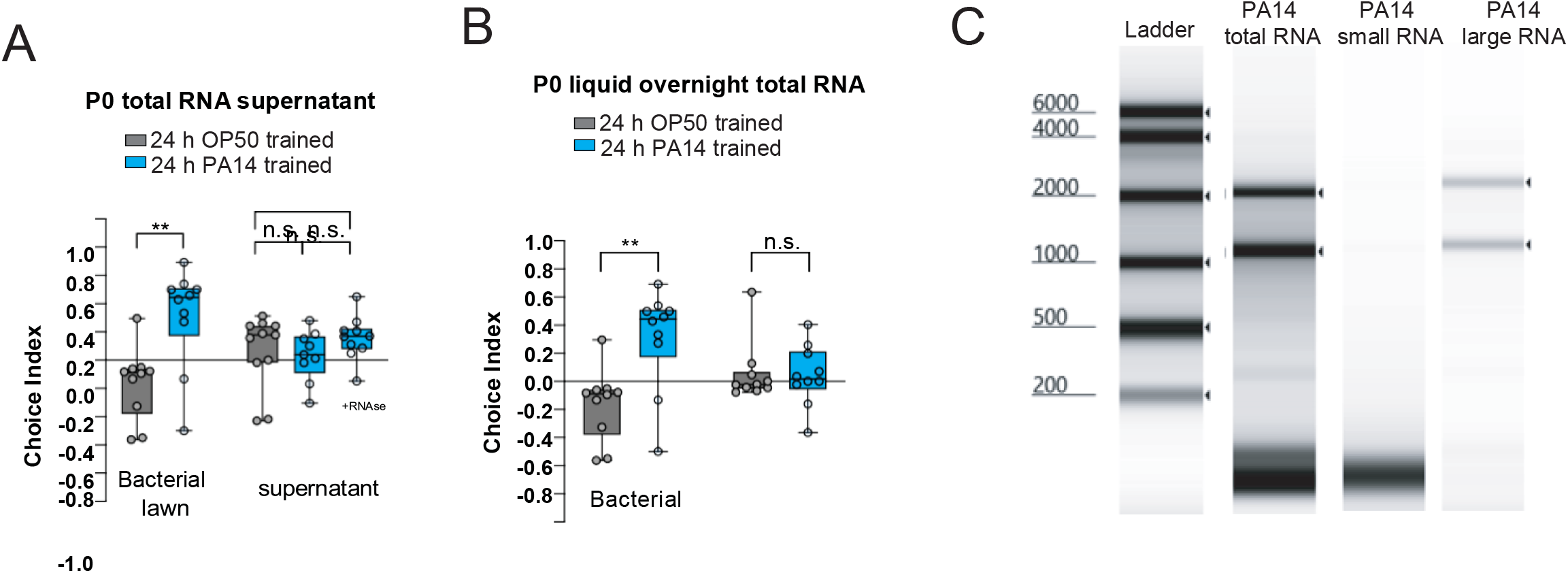
PA14 metabolites or RNA isolated from planktonic PA14 is not sufficient to induce maternal PA14 avoidance. (A) PA14 supernatant does not induce maternal avoidance of PA14. (B) Total RNA isolated from planktonic (i.e., overnight culture) of PA14 is not sufficient for maternal avoidace of PA14. One-Way ANOVA, Tukey’s multiple comparison test, mean ± SEM. n ≥ 6-10 choice assay plates with 50-200 worms per plate. For comparisons between multiple training conditions and genotypes: Two-Way ANOVA, Tukey’s multiple comparison test, mean ± SEM. n ≥ 6-10 choice assay plates with 50-200 worms. At least two biological replicates were performed. (C) Bioanalyzer results of isolated and purified PA14 total, small, and large RNAs.

**Supplemental Figure 2:**
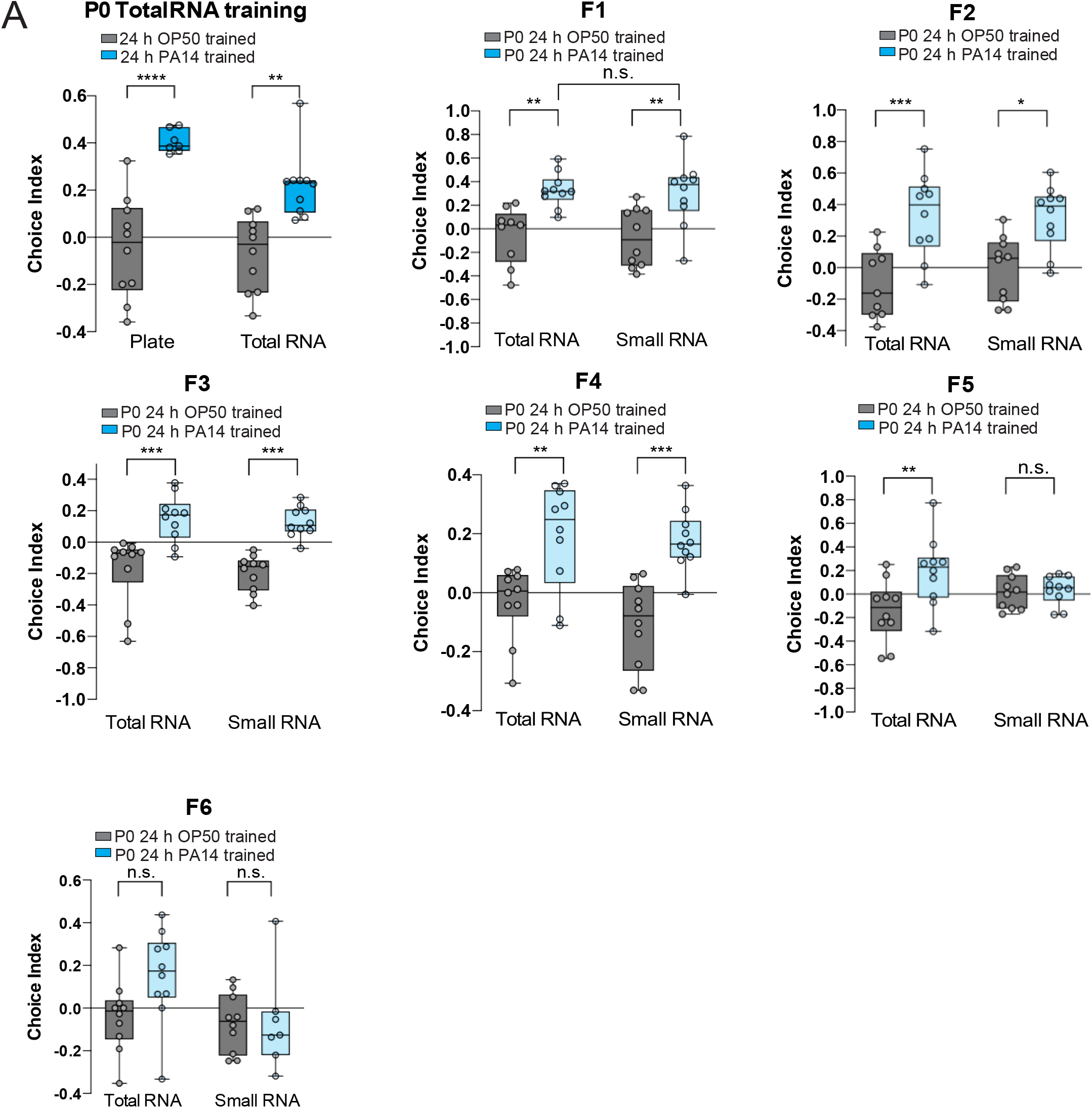
PA14 metabolites or RNA isolated from planktonic PA14 is not sufficient to induce maternal PA14 avoidance. (A) PA14 supernatant does not induce maternal avoidance of PA14. (B)_ Total RNA isolated from planktonic (i.e., overnight culture) of PA14 is not sufficient for maternal avoidace of PA14. One-Way ANOVA, Tukey’s multiple comparison test, mean ± SEM. n ≥ 6-10 choice assay plates with 50-200 worms per plate. For comparisons between multiple training conditions and genotypes: TwoWay ANOVA, Tukey’s multiple comparison test, mean ± SEM. n ≥ 6-10 choice assay plates with 50-200 worms. At least two biological replicates were performed.

**Supplemental Figure 3:**
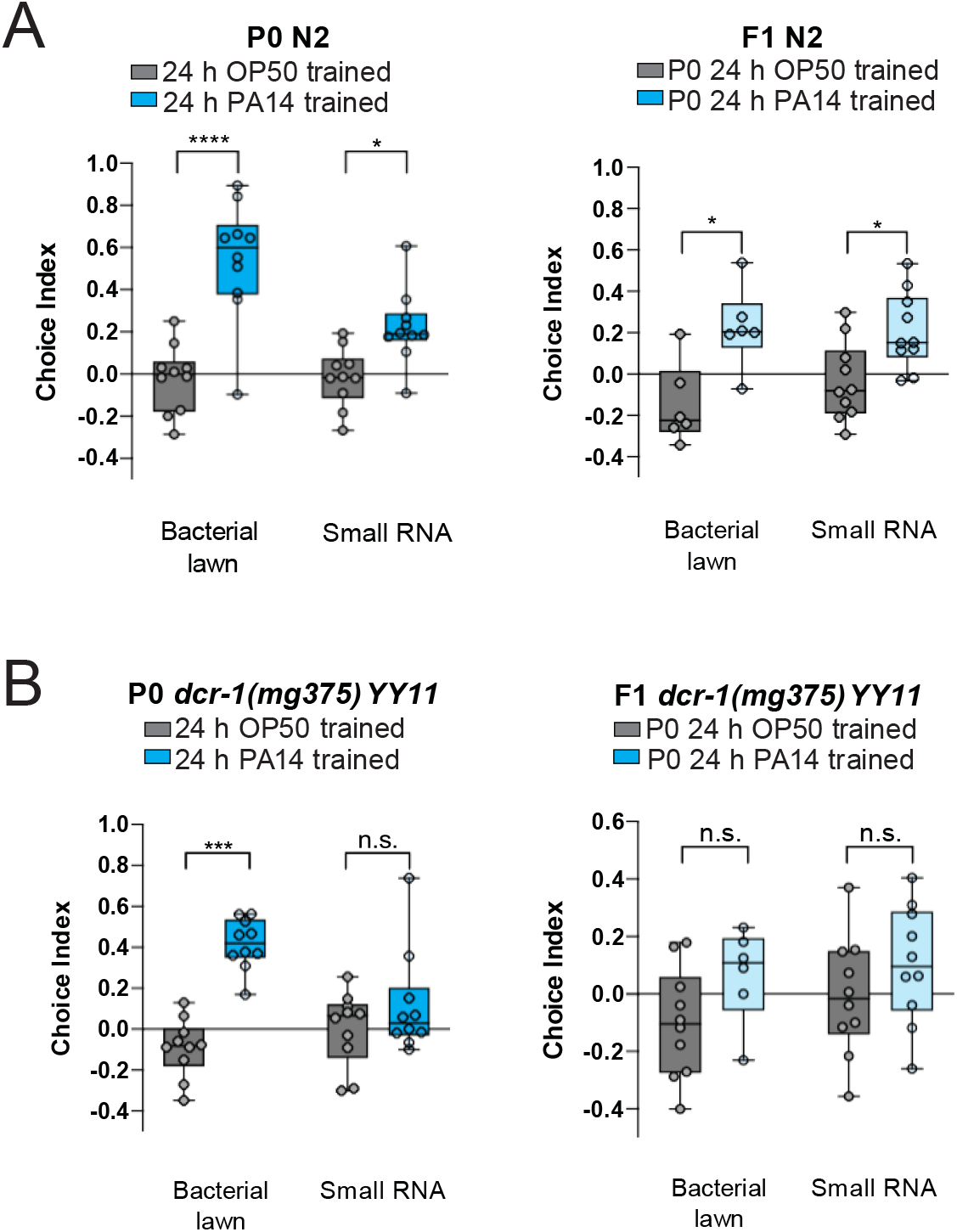
Worm dsRNA processing machinery are required for bacterial sRNA-induced pathogen avoidance. **(A)** Wild-type control for Figure 4. (B) *dcr-1(mg375)* with background *mut-16* mutation is required for PA14 small RNA induced PA14 maternal avoidance. One-Way ANOVA, Tukey’s multiple comparison test, mean ± SEM. n ≥ 6-10 choice assay plates with 50-200 worms per plate. For comparisons between multiple training conditions and genotypes: Two-Way ANOVA, Tukey’s multiple comparison test, mean ± SEM. n ≥ 6-10 choice assay plates with 50-200 worms. At least two biological replicates were performed.

**Supplemental Figure 4:**
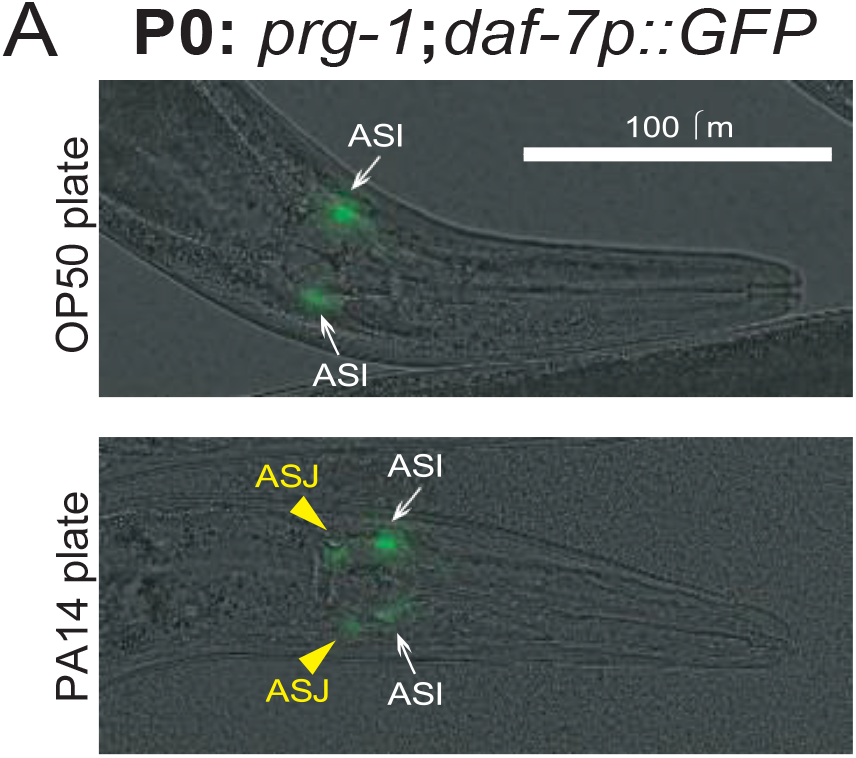
*daf-7* expression in the ASI neurons is not induced in *prg-1 mutants* upon PA14 exposure. (A) *prg-1* mutants, like wild-type worms (Fig. 4), induce *daf-7::gfp* expression in the ASJ neuron, but not the ASI neuron, after PA14 bacterial lawn exposure.

**Supplemental Figure 5:**
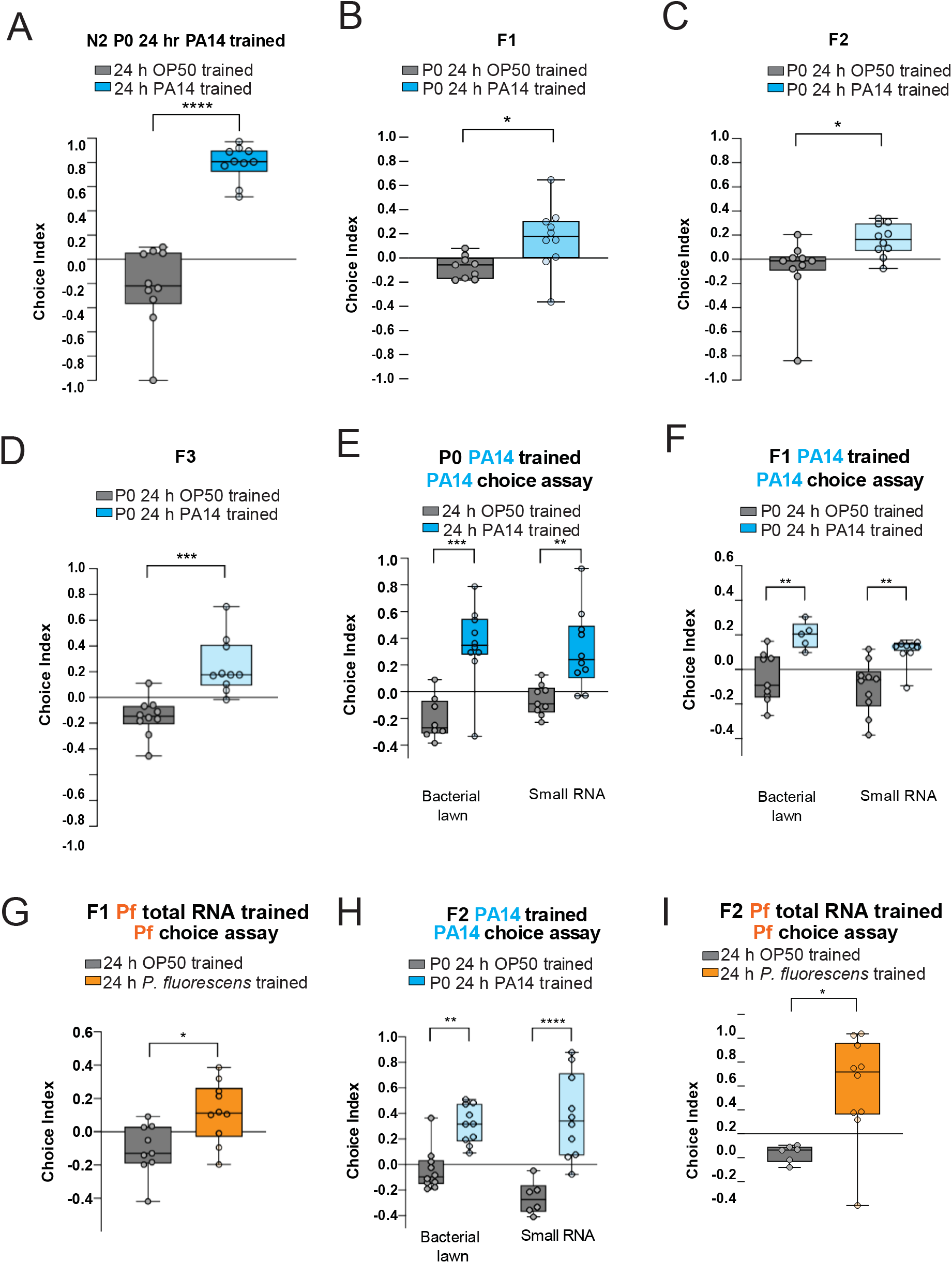
*C. elegans* uses small RNA to transgenerationally avoid environmentally abundant and pathogenic bacteria. (A-D) Untrained (naïve) progeny of PA14 bacterial lawn-trained mothers avoid PA14 in generation F1 (B), F2 (C), F3 (D). Students t-test, mean ± SEM. n ≥ 6-10 choice assay plates with 50-200 worms per plate. At least two biological replicates were performed. (E, F, H) Untrained (naïve) progeny of PA14 bacterial lawn-trained mothers (E) avoid PA14 in generation F1 (F) and F2 (H). (G, I) Total RNA from *P. fluorescens* is sufficient to induce transgenerational avoidance of *P. fluorescens* in F1 (G), and F2 (I) progeny. One-Way ANOVA, Tukey’s multiple comparison test, mean ± SEM. n ≥ 6-10 choice assay plates with 50-200 worms per plate. For comparisons between multiple training conditions and genotypes: Two-Way ANOVA, Tukey’s multiple comparison test, mean ± SEM. n ≥ 6-10 choice assay plates with 50-200 worms. At least two biological replicates were performed.

**Supplemental Table 1: DESeq2 results comparing annotated small RNAs isolated from PA14 bacteria grown at 25C or 15C on plates.**

**Supplemental Table 2: DESeq2 results comparing annotated small RNAs isolated from PA14 bacteria grown at 25C on plated or in liquid.**

**Supplemental Table 3: DESeq2 results comparing antisense coding region reads from PA14 bacteria grown at 25C or 15C on plates.**

**Supplemental Table 4: DESeq2 results comparing antisense coding region reads from PA14 bacteria grown at 25C plates or in liquid.**

**Supplemental Table 5: DESeq2 results comparing 100 bp windows of sense and antisense reads from PA14 small RNAs isolated from bacteria grown at 25C or 15C on plates.**

**Supplemental Table 6: DESeq2 results comparing sense and antisense reads from PA14 small RNAs isolated from bacteria grown at 25C plates or in liquid.**

